# Space wandering in the rodent default mode network

**DOI:** 10.1101/2023.08.31.555793

**Authors:** Trang-Anh Estelle Nghiem, Byeongwook Lee, Tzu-Hao Harry Chao, Nicholas K. Branigan, Percy K. Mistry, Yen-Yu Ian Shih, Vinod Menon

## Abstract

The default mode network (DMN) is a large-scale brain network known to be suppressed during a wide range of cognitive tasks. However, our comprehension of its role in naturalistic and unconstrained behaviors has remained elusive because most research on the DMN has been conducted within the restrictive confines of MRI scanners. Here we use multisite GCaMP fiber photometry with simultaneous videography to probe DMN function in awake, freely exploring rats. We examined neural dynamics in three core DMN nodes— the retrosplenial cortex, cingulate cortex, and prelimbic cortex— as well as the anterior insula node of the salience network, and their association with the rats’ spatial exploration behaviors. We found that DMN nodes displayed a hierarchical functional organization during spatial exploration, characterized by stronger coupling with each other than with the anterior insula. Crucially, these DMN nodes encoded the kinematics of spatial exploration, including linear and angular velocity. Additionally, we identified latent brain states that encoded distinct patterns of time-varying exploration behaviors and discovered that higher linear velocity was associated with enhanced DMN activity, heightened synchronization among DMN nodes, and increased anticorrelation between the DMN and anterior insula. Our findings highlight the involvement of the DMN in collectively and dynamically encoding spatial exploration in a real-world setting. Our findings challenge the notion that the DMN is primarily a “ task-negative” network disengaged from the external world. By illuminating the DMN’s role in naturalistic behaviors, our study underscores the importance of investigating brain network function in ecologically valid contexts.

**Significance statement:** Our research advances understanding of the default mode network (DMN), a brain network implicated in numerous neuropsychiatric and neurological disorders. In contrast to previous research examining immobilized subjects, we took the novel approach of investigating DMN function during naturalistic behaviors in freely moving rodents. Using a combination of multisite fiber photometry, video tracking, and computational modeling, we discovered a prominent role for the DMN in naturalistic real-world spatial exploration. Our findings challenge conventional views that the DMN is disengaged from interactions with the external world and underscore the importance of probing brain function in ecologically relevant settings. This work enriches our understanding of brain function and has important implications for pre-clinical investigations of disorders involving DMN dysfunction.

## Introduction

The discovery of the default mode network (DMN)—an ensemble of interconnected, distributed brain regions—has significantly advanced our understanding of the brain’s functional organization (1-5). Intriguingly, the DMN exhibits a pattern of neural activity unique among major brain networks: its responses are typically suppressed during engagement with external stimuli across cognitive tasks as diverse as working memory, selective attention, and response inhibition (4-9). This characteristic pattern, along with the DMN’s elevated “ resting-state” activity (10-13), and suppression during cognitively demanding tasks has led some researchers to consider it a “ task-negative” system that is detached from the external world (14, 15). Such a characterization is reinforced by observations linking inadequate DMN suppression to attentional lapses and compromised cognitive performance (8, 16, 17). However, while its “ task-negative” suppression shaped our initial understanding of the DMN, a growing body of evidence is broadening our conception of its functions (5). Far from being solely disengaged during active cognition, the DMN may be integral to a variety of complex, internally oriented, mental processes such as mind wandering, autobiographical memory, and planning of future events (3, 5, 18, 19). This dichotomy between the DMN’s suppression during external-stimulus-driven cognition and its hypothesized role in internally focused mental processes remains enigmatic. Specifically, it is not known whether the DMN is engaged during internally driven, self-motivated interaction with the external world. A comprehensive understanding of the DMN’s contribution to cognition, particularly in the context of naturalistic behaviors and environments, has remained elusive. Here we seek to elucidate this understudied aspect of DMN functionality and evaluate a richer picture of its contribution to brain function under ecologically valid conditions.

Most of our current understanding of the DMN comes from non-invasive functional magnetic resonance imaging (fMRI) studies (5). While this methodology has provided invaluable insights, it imposes significant constraints: participants must remain immobile, which precludes the study of naturally occurring, unrestrained behaviors. Moreover, the temporal resolution of fMRI is limited, capturing only slower hemodynamic responses rather than the fast-changing neural dynamics that may be more directly tied to ongoing behaviors (20, 21). These limitations not only limit our understanding of the DMN’s involvement in naturalistic settings but also leave an important gap in our comprehensive understanding of this crucial brain network’s roles in dynamic, real-world scenarios.

Invasive neuronal recordings in rodents offer a promising avenue to overcome the limitations inherent to fMRI studies. The identification of a rodent analog to the human DMN presents a unique opportunity to extend our understanding of this network’s role in naturalistic settings (22-28). The rodent DMN has primarily been investigated through functional connectivity analysis of resting-state fMRI data acquired while the animal is immobile under anesthesia (22, 25-27, 29-32). Key cortical nodes in the rodent DMN, such as the retrosplenial cortex (RSC), cingulate cortex (Cg), and prelimbic cortex (PrL), have been identified primarily through these methodologies. Similar to its human counterpart, the rodent DMN shows strong synchronized activity between the medial parietal cortex and medial frontal cortex (33, 34). While this parallel offers a promising basis for comparative studies, the functional role of the rodent DMN in cognition remains elusive. Previous research using invasive neuronal recordings in individual DMN regions, such as the RSC, suggests a role in spatial information processing (35-39, 40). However, these studies have been largely restricted to examining single DMN nodes in isolation. This focus on isolated regions leaves a significant gap in our understanding of how the DMN collectively encodes and orchestrates complex behaviors like spatial exploration. Our study aims to bridge this gap by employing invasive neuronal recordings across multiple nodes of the rodent DMN during free exploration, thereby providing a more comprehensive picture of the network’s role in naturalistic behaviors.

To investigate the functional role of the DMN in naturalistic behaviors, we simultaneously recorded GCaMP neural activity across multiple DMN nodes in awake, freely moving Thy1-GCaMP6f transgenic rats. We employed state-of-the-art spectrally resolved fiber-photometry techniques (24, 41, 42) to capture neural activity in three key nodes of the DMN: the RSC, Cg, and PrL nodes. We also concurrently recorded neural activity in the anterior insula (AI), a central node in the salience network (SN) known for its role in detecting important external events and adaptively guiding attention and behavior (43, 44). In both humans and rodents, attention-demanding tasks typically lead to the suppression of DMN activity while activating the AI (23, 24, 43, 45). Our selection of the specific DMN nodes was therefore based on previous studies that have demonstrated their suppression in response to both salient environmental stimuli (24) (**Figure 1B, C**) and direct optogenetic stimulation of the AI (23, 24, 45). We combined fiber-photometry, videography, and computational modeling to characterize neural dynamics of the DMN and AI nodes during free exploration, evaluate whether DMN nodes encode behaviorally relevant information during internally driven spatial exploration, and determine whether the encoding of spatial exploration differs between DMN nodes and the AI.

**Figure 1.**
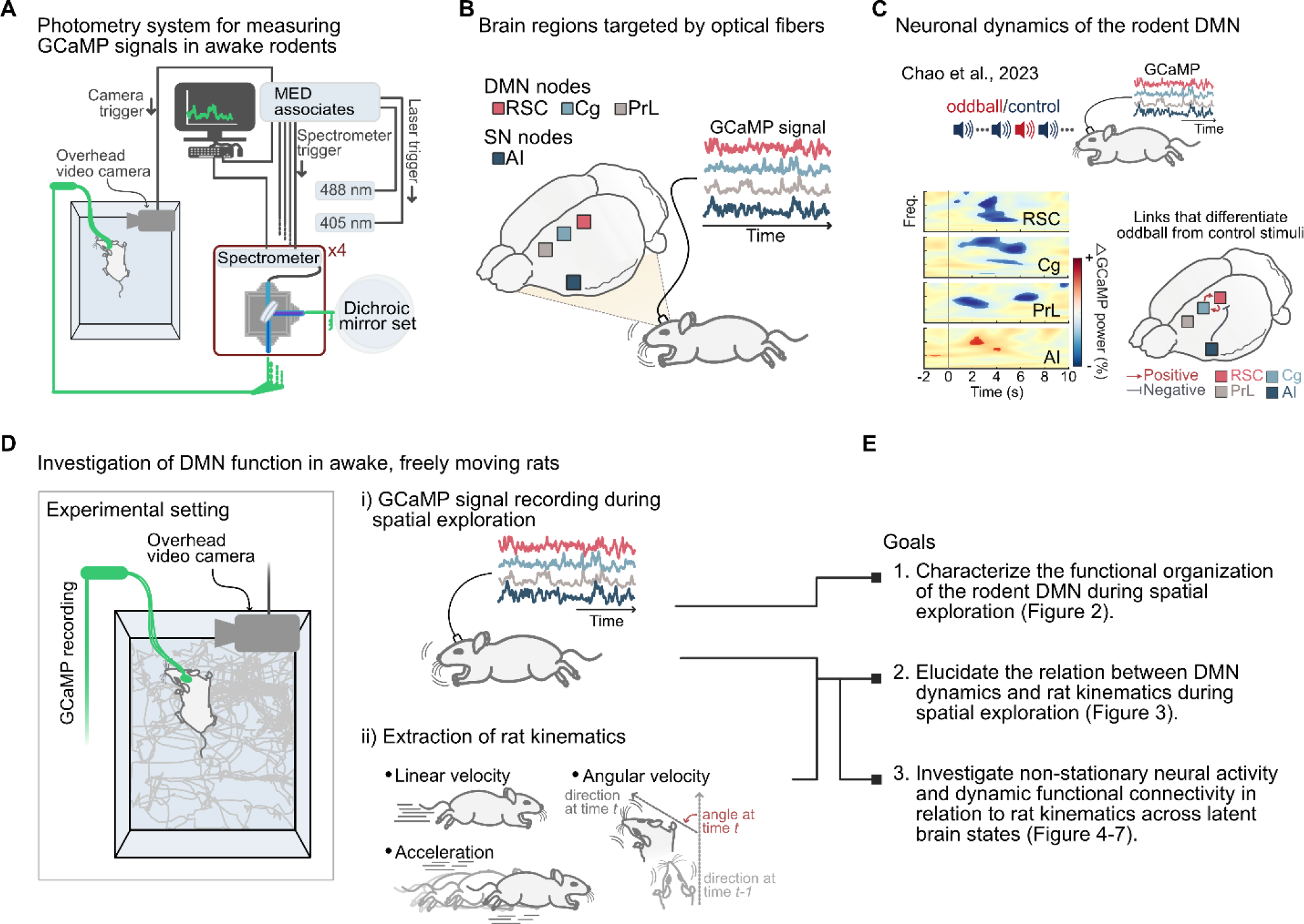
Overview of experimental design and data analysis strategy. **(A)** Schematic illustration of the fiber-photometry platform for recording neuronal GCaMP activity in awake, freely-moving rats. **(B)** Brain regions targeted by optical fibers. We recorded neuronal GCaMP activity concurrently in the RSC, Cg, PrL, and AI in each rat. **(C)** Deactivation of the rodent DMN and activation of the AI in response to salient auditory stimuli (24). Experimental design and auditory stimulation protocol (top). Time-frequency analysis of oddball-stimulus-evoked neural responses in the RSC, Cg, PrL, and AI (bottom left). Directed neural interactions in response to oddball, compared to control, stimuli (bottom right). (**D**) Overall approach and study objectives. Schematic of the freely moving rat within its environment with concurrent recording of GCaMP signals and videography (left). An example of a rat’s locomotion trajectory (in grey). Simultaneous multi-site GCaMP signals (middle top) and rodent kinematic measures extracted from videography, including linear velocity, angular velocity, and acceleration (middle bottom). **(E)** Major research goals of the current study.

We had three main goals in this study. Our first goal was to characterize the functional organization of the rodent DMN during spatial exploration, leveraging multichannel GCaMP neural recordings to test the hypothesis that the function of the DMN goes beyond conventional task-negative disengagement in ecologically valid environments. While previous studies have identified the RSC, Cg, and PrL as key components of the rodent DMN (23-28), most have used resting-state fMRI recordings under anesthesia, limiting their relevance to awake behaving states. Furthermore, while electrophysiological investigations have shed light on the RSC’s role in spatial navigation (35-37, 46-48), they have largely concentrated on this single DMN node, leaving a gap in our understanding of the broader network interactions. We used hierarchical clustering to test the hypothesis that the DMN nodes would be strongly coupled as a network distinct from the AI during free spatial exploration, and probed phase relationships and directionality of signaling among DMN nodes and the AI (24, 41, 42).

Our second goal was to investigate the relationship between temporal dynamics of DMN neural activity and behavior during internally driven spatial exploration. To achieve this, we concurrently monitored each animal’s position using videography and computed kinematic metrics, including linear velocity, angular velocity, and linear acceleration (**Figure 1D**). By probing the relationship between these kinematic measures and GCaMP neural activity, we sought to uncover whether the DMN encodes active behaviors within a naturalistic environment. We tested the hypothesis that the very DMN nodes previously shown to be suppressed during exposure to salient environmental stimuli would also encode internally driven spatial exploration.

Our final goal was to investigate dynamic patterns of neural activity and functional connectivity across the DMN and AI nodes, and to determine whether nonstationary neural patterns reflected time-varying changes in the kinematics of spatial exploration. To accomplish this, we modeled GCaMP activity with a Bayesian switching linear dynamical system (BSDS) (49). This approach allowed us to identify latent brain states, each characterized by distinct profiles of neural activation and connectivity (49). Switching linear dynamical system models (50) like BSDS provide an effective way of revealing the underlying structure and dynamics in complex time series, and these models are increasingly being used to decode meaningful latent variables underlying neuronal and fMRI recordings (23, 24, 49, 51-54). We hypothesized that BSDS modeling would reveal latent brain states corresponding to distinct kinematic patterns, thereby elucidating how spatial exploration behaviors evolve in tandem with the dynamics of neural activity and connectivity.

Our findings uncover a dynamic role for the DMN in orchestrating naturalistic, internally driven interactions with the environment, and provide a more comprehensive understanding of this pivotal network. Our results not only challenge the traditional notion of the DMN as a ‘task-negative’ system disengaged from external stimuli but also shed new light on its functionality in more ecologically relevant contexts.

## Results

### The RSC, Cg, and PrL emerge as a network distinct from the AI during spatial exploration

We utilized multisite neuronal GCaMP time series data to investigate functional connectivity between the RSC, Cg, PrL, and AI during spatial exploration. Each rat was allowed to freely explore its home cage under an awake and freely moving condition, without exposure to additional external stimuli. We detected significant functional connectivity between the DMN nodes: RSC-Cg (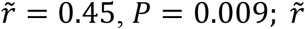 is the median Pearson correlation *r* over the 8 rats), RSC-PrL 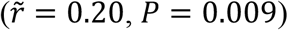, and Cg-PrL 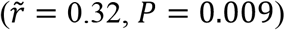 (all two-sided Wilcoxon signed-rank tests, false discovery rate (FDR) corrected) (**Figure 2A**). The RSC and Cg showed stronger functional connectivity than the RSC and PrL and the Cg and PrL (both *P* = 0.017, FDR corrected). Interestingly, DMN nodes and the AI did not display negative correlations, rather the correlations were weakly positive but significantly above zero for RSC-AI 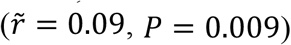 and PrL-AI 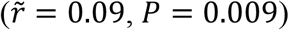 region pairs (both FDR corrected). However, functional connectivity within DMN nodes was significantly enhanced over connectivity between the DMN nodes and the AI. RSC-Cg correlations were greater than RSC-AI, Cg-AI, and PrL-AI correlations (all *P* = 0.017), RSC-PrL correlations were greater than RSC-AI and Cg-AI correlations (both *P* = 0.035), and Cg-PrL correlations were greater than RSC-AI and Cg-AI correlations (both *P* = 0.017, all FDR corrected). Overall, intra-DMN connectivity was stronger than DMN-AI connectivity, consistent with the hypothesis that the three rodent DMN nodes remain a tightly coupled network during spatial exploration.

**Figure 2.**
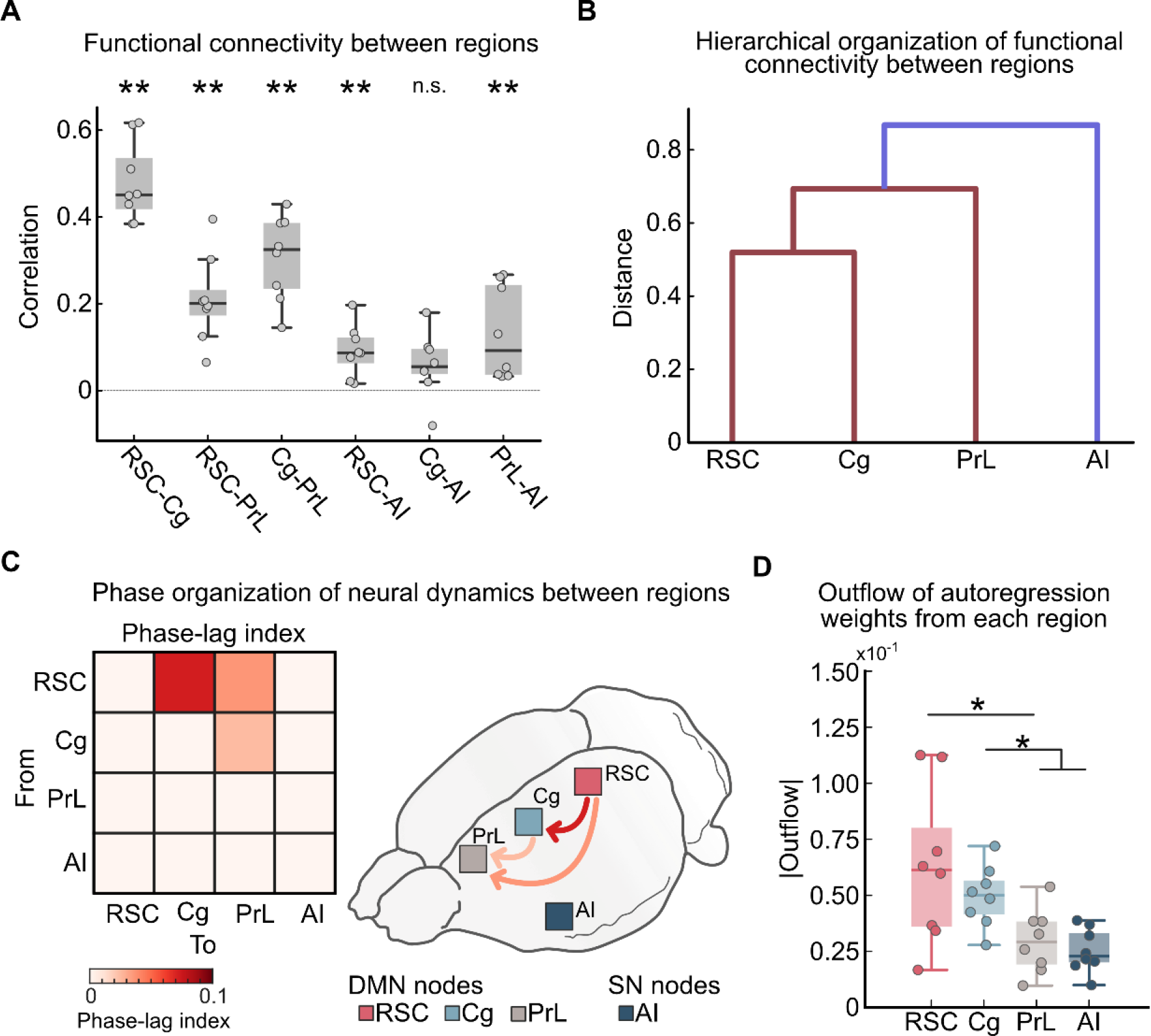
Functional interactions between DMN nodes and AI during spatial exploration. **(A)** Functional connectivity between RSC, Cg, PrL, and AI assessed using Pearson correlation (** FDR corrected *P* < 0.01, n.s. FDR corrected *P* ≥ 0.05). **(B)** Dendrogram based on hierarchical clustering of functional connectivity between the RSC, Cg, PrL, and AI. **(C)** Phase relations between the RSC, Cg, PrL, and AI measured using phase-lag index. Heat map of phase-lag index between regions, showing links that are significantly different from zero (left, thresholded at FDR corrected *P* < 0.05). Direction and strength of phase lag between RSC, Cg, PrL, and AI (right). Neural activity in the RSC led all other DMN nodes. **(D)** Directed weighted outflow from the RSC, Cg, PrL, and AI estimated using a multivariate vector autoregressive model. The RSC and Cg showed enhanced outflow compared to other regions (* FDR corrected *P* < 0.05). Circles in **A** and **D** represent data of individual rats. Box and whisker plots show 25^th^ and 75^th^ percentiles, and horizontal lines represent median values.

We then used hierarchical clustering to evaluate the relative strength of functional connectivity among the four regions. The RSC and Cg showed the strongest connectivity, while the AI showed the weakest connectivity with the three other regions, such that the RSC, Cg, and PrL formed a hierarchical cluster distinct from the AI (**Figure 2B**). Together, these results suggest a hierarchical functional organization of brain dynamics during free exploration, with functional connectivity differentiating the rodent DMN nodes from the AI.

### Neural activity in the RSC leads activity of other DMN nodes during spatial exploration

Next, we examined phase relations between the RSC, Cg, PrL, and AI. We used directed phase-lag index (PLI) (55, 56) to determine the phase relationships between neuronal GCaMP signals in each node. PLI quantifies how consistently one signal leads or lags another: a positive PLI between signal *i* and signal *j* denotes that *i* leads *j* in phase, while a negative PLI denotes that *i* lags *j* in phase. A zero PLI indicates that *i* and *j* are either perfectly synchronous or do not present consistent phase relations throughout time. PLI analysis revealed that the RSC significantly phase-led the Cg (*P* = 0.023) and the PrL (*P* = 0.023), and that the Cg significantly phase-led the PrL (*P* = 0.023) (all FDR corrected) (**Figure 2C**). No significant phase lag was observed between the RSC and AI (*P* = 0.546), Cg and AI (*P* = 0.312), and PrL and AI (*P* = 0.312) (all FDR corrected). These results demonstrate that neural activity in the RSC leads the two other DMN nodes, but not the AI, in phase, during spatial exploration.

### The RSC and Cg are outflow hubs during spatial exploration

To further investigate the directional information flows between DMN nodes and the AI, we used a multivariate vector autoregression (MVAR) model. MVAR jointly estimates the influence of activity at each node at time *t* on all other nodes at time *t* + 1. The MVAR-derived directed connectivity matrix was then used to compute net outflow from each region. This analysis revealed a higher degree of outflow from RSC compared to PrL (*P* = 0.031), from Cg compared to PrL (*P* = 0.023), and from Cg compared to AI (*P* = 0.023) (all FDR corrected) (**Figure 2D**). No significant outflow differences were found between RSC and Cg (*P* = 0.375), RSC and AI (*P* = 0.059), and PrL and AI (*P* = 0.945) (all FDR corrected). These results suggest that the RSC and Cg may serve as outflow hubs predicting dynamics in other nodes during spatial exploration.

### Neural activity in the DMN nodes is correlated with kinematics during spatial exploration

Next, we sought to determine whether neuronal activity of DMN nodes and the AI was correlated with kinematics of spatial exploration (**Figure 3**). Videographic images were used to compute three kinematic measures of spatial exploration over time: linear velocity, angular velocity, and acceleration. Linear velocity was correlated with GCaMP activity in the RSC 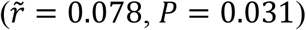, but not the Cg 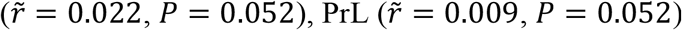, or 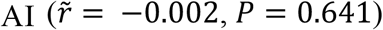 (all FDR corrected) (**Figure 3**). Angular velocity was negatively correlated with all three DMN nodes: RSC 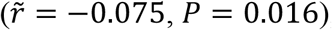, Cg 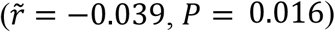, and PrL 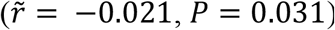, but not with the AI 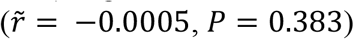 (all FDR corrected) (**Figure 3**). Acceleration was not correlated with the RSC 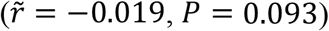, Cg 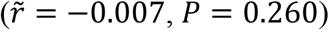, PrL, 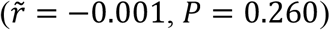, or AI 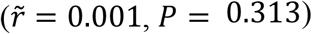 (all FDR corrected) (**Figure 3**). These results demonstrate preferential encoding of kinematics in all three nodes of the DMN, with the RSC showing the strongest and most consistent effects.

**Figure 3.**
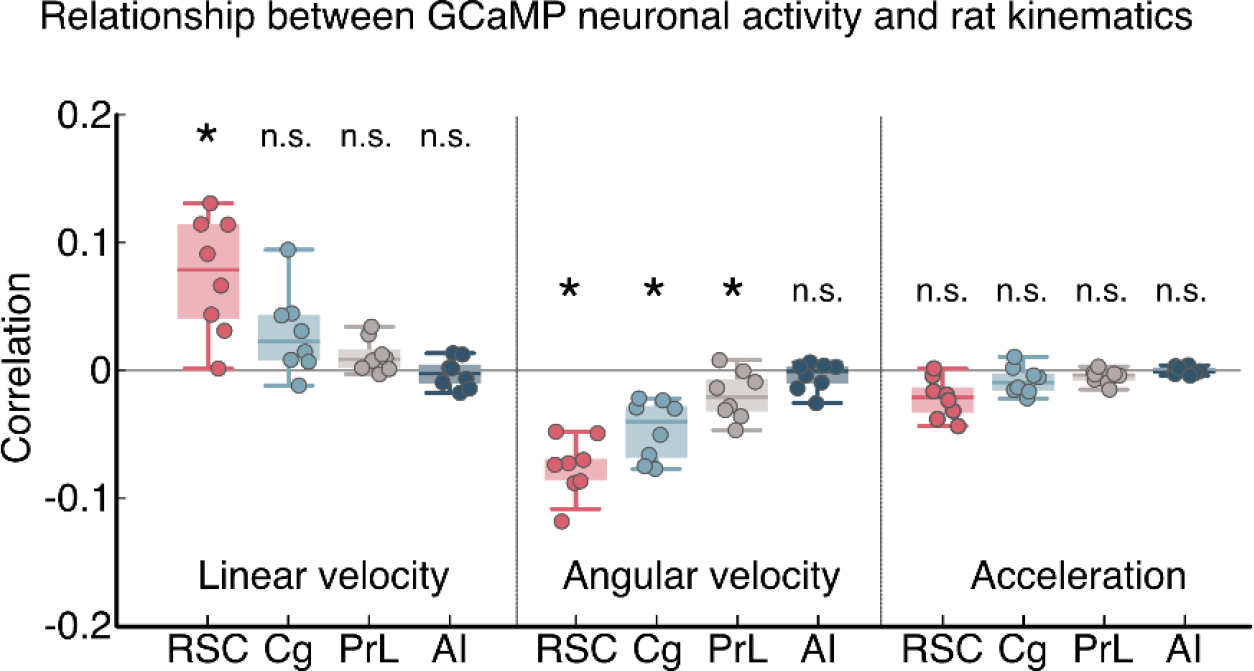
Relationship between DMN activity and kinematic measures during spatial exploration. A significant positive correlation was detected between GCaMP neural activity in the RSC and linear velocity (* FDR corrected *P* < 0.05, n.s. FDR corrected *P* ≥ 0.05). A significant negative correlation was detected between GCaMP neural activity in the RSC, Cg and PrL, and angular velocity. No correlation was found between GCaMP activity and acceleration. Results demonstrate preferential encoding of kinematics in all three nodes of the DMN, but not the AI. The RSC showed the strongest and most consistent effects. Circles represent data of individual rats. Box and whisker plots show 25^th^ and 75^th^ percentiles, and horizontal lines represent median values.

### State-space modeling reveals nonstationary neural dynamics characterized by distinct brain states

While the linear (Pearson) correlation between neural activity within the DMN and behavioral kinematics is significant, the effect sizes are weak – possibly due to the nonstationary of neural dynamics through time.. To investigate this, we first used state-space modeling to identify latent brain states based on dynamic patterns of neural activity and functional connectivity across the DMN and AI nodes. This was accomplished by applying BSDS (49), a state-space model that uncovers latent brain states between which the brain dynamics transition back and forth throughout time (**Figure 4A)**. BSDS identified 7 distinct latent brain states (**Figure 4B)**. These states varied in occupancy rate, the fraction of time that a state is inferred to occur, from state 1 with the highest (mean (SD) = 17.6% (2%)) to state 7 with the lowest (mean (SD) = 9.6% (1%)) (**Figure 4C**). These results reveal nonstationary neural dynamics associated with the DMN and AI during spatial exploration.

**Figure 4.**
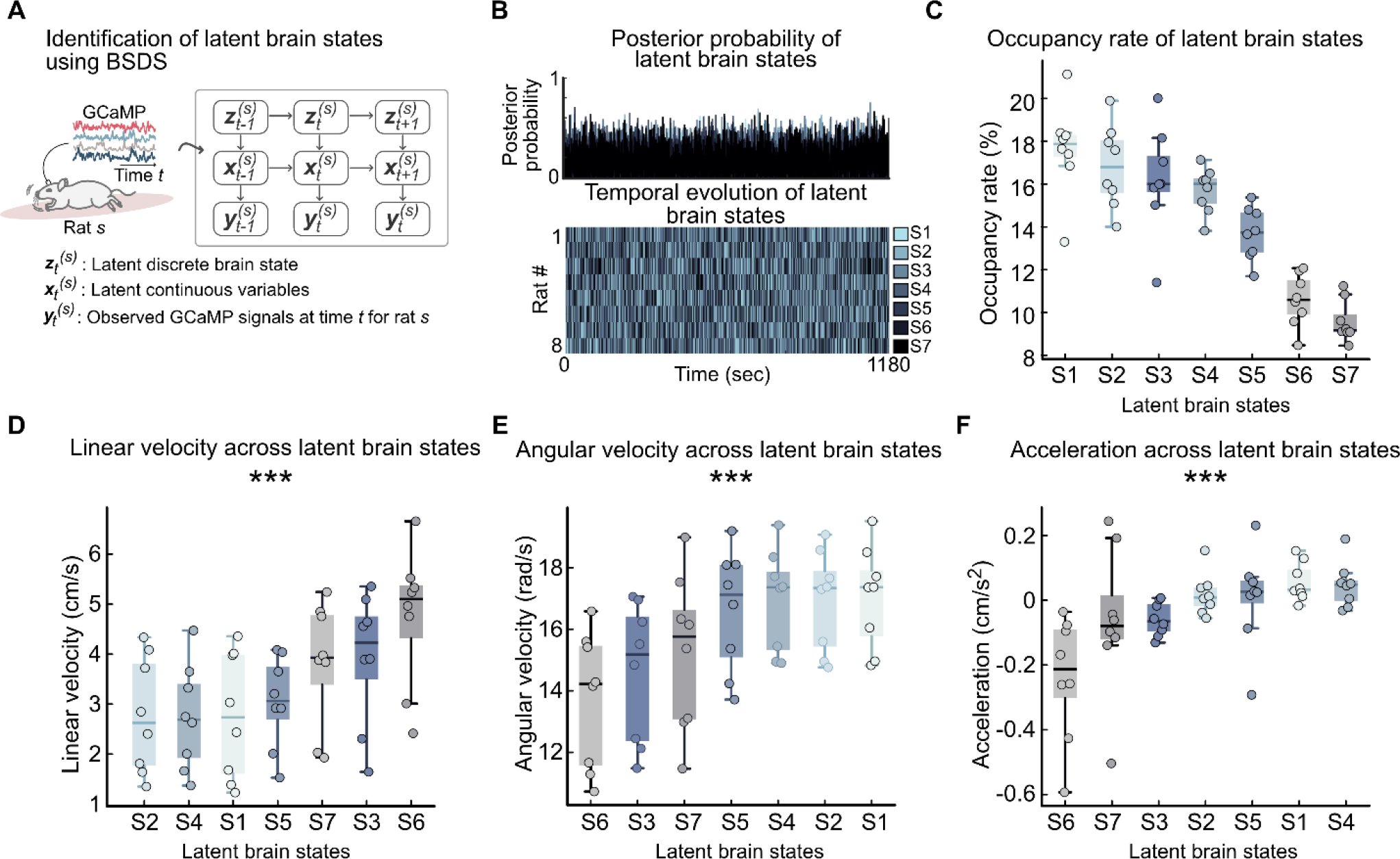
Dynamic brain states encode time-varying kinematics during spatial exploration. **(A)** Schematic of the state-space model used to obtain latent discrete brain states from the neuronal GCaMP signals. **(B)** Temporal evolution of the posterior probability of latent brain states (top) and most likely sequences of brain states in each of the eight rats (bottom). **(C)** Occupancy rate of latent brain states, ranked from the most to least frequently occupied. **(D-F)** Linear velocity, angular velocity, and acceleration vary significantly across latent brain states (*** *P* < 0.001).

### Latent brain states encode time-varying kinematics during spatial exploration

Next, we determined whether nonstationary patterns reflected time-varying changes in the kinematics of spatial exploration. Specifically, we evaluated whether latent brain states identified by BSDS were associated with different patterns of spatial exploration behaviors. We found that the brain states differed in linear velocity (*P* < 0.001), angular velocity (*P* < 0.001), and acceleration (*P* < 0.001) (**Figure 4D-F**) (all Friedman tests). These results indicate that the latent brain states are behaviorally relevant, and that they encode distinct nonstationary patterns of kinematics during spatial exploration.

### Neural activity in the DMN encodes time-varying kinematics during spatial exploration

We extended this analysis to examine how neural activity in the DMN nodes and AI covaried with time-varying kinematics across the seven latent brain states. Brain states with higher neural activity exhibited higher linear velocity (**Figure 5A**). Specifically, mean GCaMP activity in the 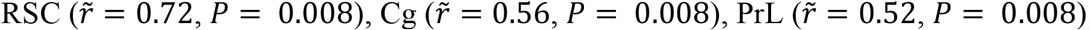, and 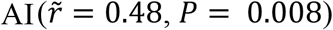 was positively correlated with linear velocity across brain states. The strength of this correlation was higher in the RSC compared to the AI (FDR corrected *P* = 0.023) (**Figure 5B**). Angular velocity and acceleration displayed a convergent pattern of correlations with neural activity in the DMN (**Supplementary Results** and **Figure S1**). These results demonstrate that neural activity in the DMN encodes time-varying kinematic information, with the strongest effects being observed in the RSC node.

**Figure 5.**
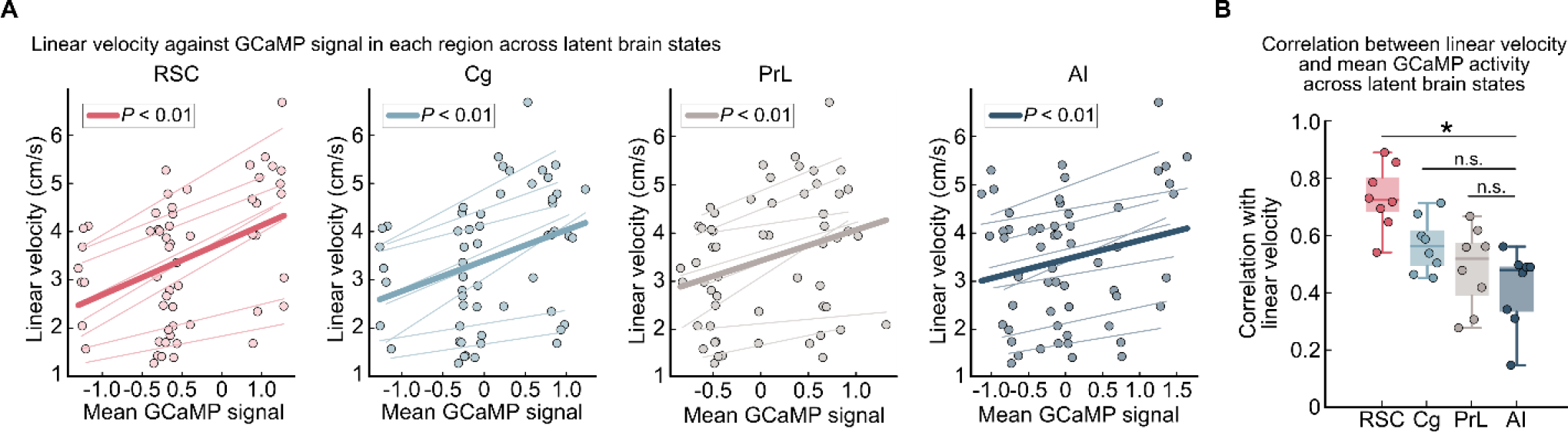
DMN activity encodes time-varying linear velocity during spatial exploration across latent brain states. **(A)** Mean linear velocity as a function of mean GCaMP signal amplitude in the RSC, Cg, PrL, and AI, across latent brain states. Linear velocity was positively correlated with GCaMP activity in all three DMN nodes as well as the AI. Lighter lines represent linear fits across latent brain states in each rat, and each circle represents data from a specific rat and latent brain state. Dark lines represent linear models across all rodents. **(B)** Strength of correlation between linear velocity and mean GCaMP signal in the RSC, Cg, PrL, and AI. The correlation was higher in the RSC than in the AI (* FDR corrected *P* < 0.05, n.s. FDR corrected *P* ≥ 0.05). Box and whisker plots show 25^th^ and 75^th^ percentiles, and horizontal lines represent median values. Dots in each figure represent data points from individual rats.

### Spectral entropy of neural activity in the DMN varies with time-varying kinematics during spatial exploration

Next, we examined how spectral entropy of GCaMP neural activity changed with time-varying kinematics across the 7 brain states. Spectral entropy is a measure of neural variability that has been linked to irregularity and complexity associated with information processing capability (56-58). Spectral entropy of GCaMP neural activity was negatively correlated with linear velocity across brain states in the RSC 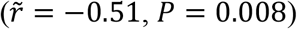, Cg 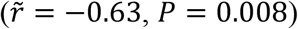, PrL 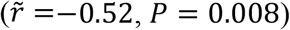, and AI 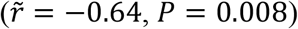 (**Figure S2A**). Angular velocity and acceleration displayed a convergent pattern of correlations with spectral entropy in the DMN and AI (**Supplementary Results** and **Figure S4**). These results demonstrate that brain states with lower spectral entropy exhibit higher linear velocity, suggesting that these states exhibit more structured dynamics and information processing. In contrast, brain states with higher angular velocity and acceleration were characterized by higher entropy, perhaps reflecting greater neural variability associated with more complex aspects of spatial exploration.

### Functional connectivity of DMN nodes encodes time-varying kinematics during spatial exploration

Finally, we evaluated how functional connectivity covaried with rat kinematics across the 7 brain states. Linear velocity was positively correlated with functional connectivity between the RSC and Cg 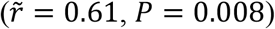 and between the Cg and PrL 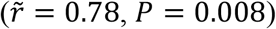 (**Figure 6**). In contrast, linear velocity was negatively correlated with functional connectivity between the RSC and AI 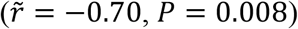. Angular velocity and acceleration also covaried significantly with functional connectivity of DMN nodes across the latent brain states (**Supplementary Results** and **Figures S3, S4**). These results demonstrate that heightened activity and synchronization within the DMN, together with anticorrelated activity between the RSC node of the DMN and the AI, encode time-varying kinematics during spatial exploration.

**Figure 6.**
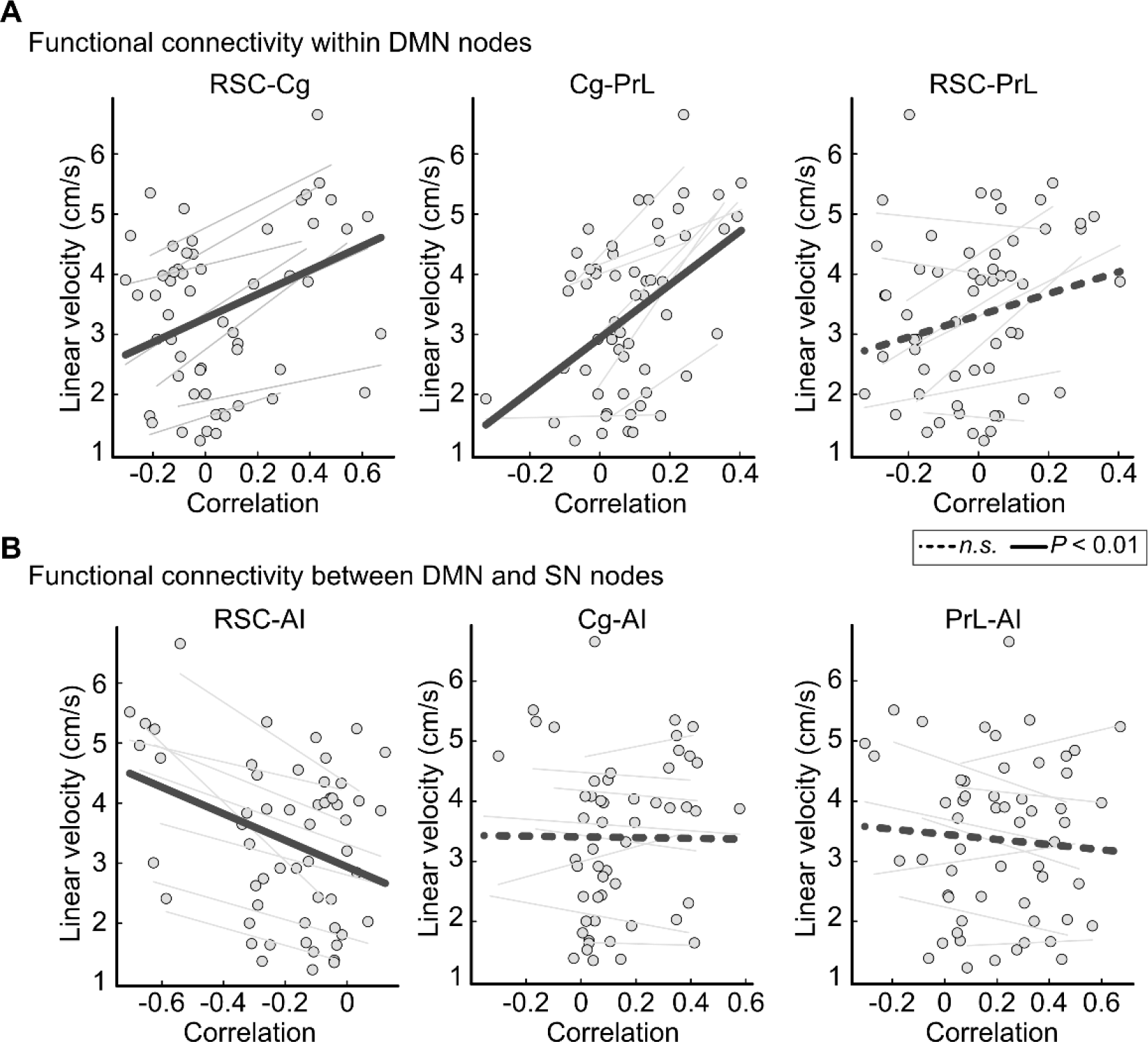
Functional connectivity among the DMN and AI encodes time-varying linear velocity during spatial exploration. **(A)** Mean linear velocity as a function of intra-DMN functional connectivity across latent brain states. Linear velocity was positively correlated with functional connectivity between the RSC and Cg, and between the Cg and PrL. **(B)** Mean linear velocity as a function of connectivity between the AI node of the salience network and the DMN nodes. Linear velocity was negatively correlated with functional connectivity between the RSC and AI. Lighter lines represent linear fits across latent brain states in each rat, and each circle represents data from a specific rat and latent brain state. Dark lines represent linear models across all rodents.

**Figure 7** summarizes key results of our state-space modeling which revealed brain states characterized by distinct patterns of nonstationary time-varying behaviors during spatial exploration.

**Figure 7.**
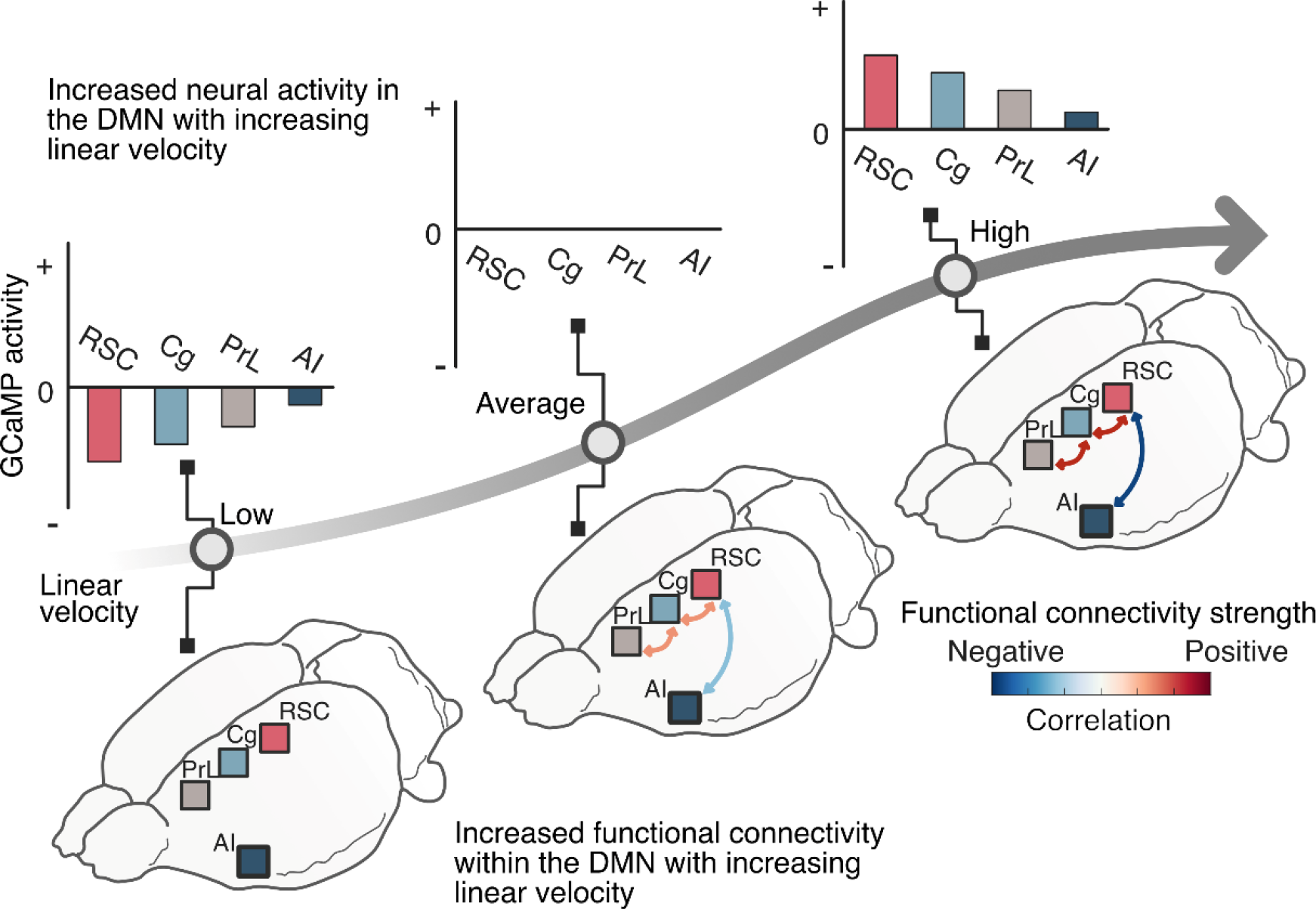
Schematic of nonstationary dynamics of the DMN and their relation to linear velocity during spatial exploration. Neural activity in the DMN and the strength of functional connectivity between DMN nodes increase with linear velocity. In contrast, the RSC and AI become more anticorrelated with linear velocity. Thus, the spatiotemporal dynamics of DMN neural activity and functional connectivity collectively encode spatial exploration.

## Discussion

Our study provides compelling evidence for a dynamic role of the DMN during internally driven spatial exploration. Even though the DMN comprises brain regions that are commonly suppressed during a wide range of tasks involving attention to external stimuli, in humans, it has also been implicated in cognitive processes like episodic memory, spatial navigation, and future planning (1-5, 59). Critically, the DMN’s function in naturalistic settings and its neural underpinning has remained elusive. Addressing this knowledge gap, we analyzed multichannel GCaMP signals from distributed brain areas and probed how three central nodes of the rodent DMN, and the AI node of the salience network, dynamically encode naturalistic, self-initiated spatial exploration behavior. Our findings revealed a hierarchical functional organization characterized by stronger neural synchronization within the DMN than between the DMN and AI during spatial exploration. In contrast to the AI, all DMN nodes encoded the kinematics of spatial exploration, with the most pronounced effects observed in the RSC. Additionally, spatial exploration was associated with distinct brain states that encoded time-varying kinematics of spatial exploration. As summarized in **Figure 7**, higher linear velocity was associated with elevated DMN activity and enhanced synchronization among DMN nodes, as well as an anticorrelation between the AI and RSC. Our findings deepen our understanding of the DMN’s neural dynamics and their pivotal role in self-initiated exploratory behaviors. This evidence challenges the view of the DMN as a predominantly ‘task negative’ network.

### Hierarchical functional organization of the DMN and AI in awake, freely moving rodents

The first goal of our study was to delineate the functional organization of the RSC, Cg, and PrL nodes of the DMN, in contrast to the AI, during active spatial exploration in awake and freely moving rodents. Prior research using fMRI has identified these regions as key cortical nodes of the rat DMN (23, 24). Crucially, however, our understanding of the DMN has been constrained by a paucity of evidence from neuronal recordings during naturalistic behaviors, and the fact that DMN investigations in rodents have been primarily performed under anesthesia—an experimental condition that amplifies low-frequency synchronization across brain regions and could potentially bias the characterization of the DMN (24). Studies utilizing resting-state fMRI data from non-anaesthetized rodents have suggested that the DMN may form a functional network separate from the salience network (22). However, as these recordings were performed in the highly confined environment of an MRI scanner, it raises questions about the functional organization of the DMN in naturalistic conditions in awake, behaving rodents.

Our multichannel GCaMP neural recordings overcome these limitations and elucidate the functional organization of DMN dynamics in freely moving rodents. Results revealed that the RSC, Cg, and PrL are more strongly synchronized with each other than with the AI. Hierarchical clustering validated this pattern and highlighted the distinction of the AI from the RSC, Cg, and PrL. This pattern of stronger synchronization within the RSC, Cg, and PrL, compared with the AI, was consistently observed in each individual rodent. Among the four nodes, the RSC and Cg demonstrated the most robust functional coupling, positioning these regions as anchors of the rodent DMN. Collectively, these results indicate that the RSC, Cg, and PrL form a distinct network separate from the AI in the awake behaving state.

Next, we investigated directed interactions between DMN nodes during spatial exploration. Specifically, we examined whether the activity of any specific DMN node preceded and predicted the activity of other nodes. Phase-lag analysis, which quantifies the temporal relationships between two neural signals by measuring their phase difference, revealed that RSC activity led activity in other DMN nodes but not the AI. Additionally, joint linear modeling of the temporal relations among the RSC, Cg, PrL and AI using a multivariate autoregressive model revealed that the RSC and Cg displayed enhanced directed outflow among all nodes. Together, results reveal organized directed relationships between DMN nodes, with the RSC preceding and predicting dynamics in the other network nodes.

In summary, these findings elucidate the hierarchical functional organization of the DMN and AI in awake, freely moving rodents, and suggest a crucial role for the RSC as a signaling hub during spatial exploration. We posit that these properties could allow the RSC to broadcast behaviorally relevant spatial information across distributed nodes of the DMN and facilitate efficient coordination of brain networks to effectively guide and modulate spatial exploration behaviors (60). Further research, employing electrophysiological recordings acquired at higher sampling rates, as well as causal manipulation of the RSC with concurrent recording in other DMN nodes, is needed to test this hypothesis.

### DMN nodes encode kinematics of spatial exploration

The second goal of our study was to investigate the interplay between dynamic DMN activity and behavioral measures associated with internally driven spatial exploration. To accomplish this, we utilized videography to compute three kinematic measures: linear velocity, angular velocity, and acceleration. Our findings reveal neural encoding of kinematics in the DMN but not in the AI.

Only the RSC encoded linear velocity, whereas all three DMN nodes, the RSC, Cg, and PrL, encoded angular velocity, with RSC being the most significantly correlated with multiple kinematic measures. The AI was the only region that did not encode either linear or angular velocity. These findings echo prior studies, which found that RSC neurons are tuned to speed (46, 59), and extend them to establish a link with angular velocity in environments free from task constraints, specific spatial layouts, and reward associations. Moreover, our findings based on GCaMP neuronal activity align with extensive human neuroimaging research demonstrating RSC involvement during spatial navigation in virtual-reality environments. Numerous human fMRI studies have revealed that RSC activity encodes distance from home (61), familiarity (62, 63), and permanence (64) of spatial landmarks and visual cues. Our study broadens these insights from controlled and virtual environments to a naturalistic setting and underscore the ecological relevance and translatability of these discoveries.

A noteworthy discovery was the broad encoding of angular velocity across DMN nodes, not limited to the RSC (**Figure 3**). This more extensive representation of angular versus linear velocity may reflect the inherent complexity of processing and decision-making associated with direction changes. Angular velocity encapsulates not just movement speed, but also the orientation and turning required to navigate an environment. Such behavior necessitates complex spatial representations, multisensory input integration, and coordinated motor responses for direction adjustment. Our results hint at a distributed representation of kinematics-related information across multiple DMN nodes, which may facilitate the intricate decision-making processes required for constrained directional changes in space.

### DMN activity and dynamical functional connectivity encode time-varying spatial exploration: insights from state-space modeling

The final goal of our study was to investigate time-varying changes in DMN activity in relation to the kinematics of spatial exploration. We employed a Bayesian dynamical system model to identify time-varying latent brain states associated with distinct patterns of neural activity, and mapped brain dynamics with behavioral changes over time. State space modeling revealed 7 distinct states in awake freely moving rats, with each state displaying different neural activity and connectivity between the DMN nodes and the AI.

Crucially, latent brain states, which were identified solely based on neural GCaMP signals, were associated with significant fluctuations in linear velocity and angular velocity. Across brain states, average neural activity in the DMN nodes increased with linear velocity and decreased with angular velocity. This relationship was significantly stronger in the RSC compared to the AI. In parallel, synchronization of DMN nodes also increased with linear velocity and decreased with angular velocity. Moreover, as linear velocity increased, spectral entropy decreased, suggesting more structured, less random neural dynamics, likely reflecting more efficient encoding of behaviorally relevant information (56, 57, 65-68).

Interestingly, the correlations between GCaMP neural activity and kinematic measurements were markedly stronger when examined across latent brain states, as compared to correlations across the full span of the recordings (compare **Figures 5** and **3**). This observation emphasizes the nonstationary nature of neural activity patterns associated with spatial exploration and underscores the value of state-space modeling. By considering the temporal evolution of latent brain states, we were able to more accurately and comprehensively characterize the relationship between neural and behavioral dynamics during spatial exploration.

Convergent with the analysis noted in the previous section, state-space modeling also revealed opposing patterns of associations between linear velocity and acceleration. This reflects physical constraints on movement as high velocity locomotion states may require deceleration to avoid obstacles and make turns. In contrast, the RSC and AI became more anticorrelated with increasing linear velocity. Shifts in functional connectivity patterns that coincide with increasing linear velocity demonstrate DMN and AI adaptability in guiding behavior during naturalistic exploration. This dynamic modulation of network activity underscores the intricate relationship between DMN function and behavior in real-world settings. The increased anticorrelation between the DMN and AI in states associated with heightened linear velocity parallels the antagonistic AI-DMN interactions during processing of salient external stimuli. These results emphasize the significant impact of locomotion kinematics in shaping state-specific neural dynamics within the rodent DMN and highlight the influence of behavioral context on network activity.

Our findings reveal dynamic reorganization of collective DMN dynamics in relation to ongoing changes in movement kinematics. Selective manipulations of DMN activity with optogenetic and chemogenetic tools are needed to further clarify the causal role of the RSC and other regions in network functions associated with spatial exploration. However, based on our observations that the RSC displayed the strongest effects of motion kinematics, and the phase-lag analysis demonstrating RSC responses as preceding responses in other DMN nodes, we suggest that the RSC may play a role in broadcasting locomotion-relevant information, thereby enabling the DMN-wide encoding of spatial exploration.

## Conclusion

Our study advances knowledge of DMN function and elucidates how its dynamics are influenced by internally driven spatial exploration in naturalistic environments. By examining GCaMP signals in awake, freely moving rats, we demonstrate that DMN activity and connectivity dynamically encode time-varying kinematics of spatial exploration. Our discovery that naturalistic behavioral patterns strongly correlate with distinct patterns of neural activity and synchronization across specific brain states emphasizes the dynamic and flexible nature of behavioral encoding by the DMN.

The research presented here expands our view of the DMN beyond resting-state synchrony and task-related deactivation. Our study reveals that the DMN, typically known for its suppression in response to external stimuli, also plays a critical role in encoding self-driven exploratory behaviors. This broadens our conceptualization of the DMN’s functionality and highlights its involvement in complex behaviors and interactions with the external world. These insights not only deepen our comprehension of the DMN’s role in cognition, but also stress the importance of studying brain network function within ecologically valid contexts to appreciate the full extent of their functional roles.

In closing, we propose that “ space wandering” —the internally driven spatial exploration behavior we have investigated here—may reflect an evolutionary precursor to mind wandering (5). Two observations make this idea tempting. First, both space and mind wandering are self-driven, arise spontaneously, and can operate independently of external goals. Second, the processes may share a neural basis in the DMN: our work suggests a DMN role in space wandering, and findings from multiple methodologies provide replicable evidence that the DMN is involved in mind wandering (5). Together, these connections suggest an evolutionary link between these complex cognitive processes across different species. This perspective may open new avenues for research into the evolutionary foundations of the DMN and bolsters the translational value of studying DMN function in awake behaving animals.

## Methods

Detailed procedures employed for data acquisition and analysis are outlined in the **Supplementary Methods**. Key steps are summarized below.

### Multisite GCaMP Recordings

Eight Thy1-GCaMP6f transgenic male Long-Evans rats were utilized to assess cortical output activity from pyramidal neurons. The rats were anesthetized, secured in a stereotactic frame, and specific coordinates in the RSC, Gg, PrL and AI were designated for the implantation of optical fibers. Spectrally resolved measurements of GCaMP signals were captured using a 4-channel fiber-photometry platform (24, 41, 42, 69-71). Subsequent to anesthetization, patch cables were attached to the implanted fibers, and the rats were allowed to recover prior to recording. The experiment was conducted within the rats’ familiar home cage environment to mitigate anxiety, stress, and extraneous environmental influences. All procedures adhered to ethical guidelines and received approval from the University of North Carolina’s Institutional Animal Care and Use Committee.

### GCaMP and Behavioral Data Analysis

#### Preprocessing

GCaMP data were preprocessed following methods detailed in a previous study (24). GCaMP signals were detrended, mean-corrected, and band-pass-filtered at 0.1 to 1.5 Hz for further analysis.

#### Functional connectivity

Pearson correlation was used to compute functional connectivity during spatial exploration between regions (RSC, Cg, PrL, and AI) taken pairwise. Hierarchical clustering was used to elucidate the overall functional organization of DMN nodes and the AI.

#### Phase-lag

Phase-lag index analysis was applied to determine phase relationships between the four brain regions.

#### Multivariate autoregression modeling

A multivariate autoregression was used to model directed regional interactions during spatial exploration by inferring linear directional weights between distinct neural recordings over time.

#### Kinematic measures associated with spatial exploration

Linear velocity, angular velocity, and acceleration were estimated as functions of time by extracting the trajectory of the rodent’s head movement derived from video recordings.

#### Relation between GCaMP neuronal activity and kinematic measures

Pearson correlation was used to evaluate the relationship between kinematic measures and GCaMP signals in each of the four brain regions.

#### State-space modeling of nonstationary neural dynamics

A Bayesian switching linear dynamical system (23, 49 {Fox, 2008 #167, 51)} was fit to time-series data to identify latent brain states associated with fluctuating neural GCaMP activity and functional connectivity patterns. Model parameters were determined using variational inference.

#### Relation between brain-state-related neural activity, spectral entropy, and functional connectivity, and time-varying kinematics

The relation between GCaMP-derived measures and kinematic measures was evaluated across latent brain states using Pearson correlation.

#### Significance testing

All significance testing was done nonparametrically. Paired-group data were submitted to two-sided Wilcoxon signed-rank tests, with FDR correction for multiple comparisons where appropriate. Data from more than two dependent groups were submitted to Friedman tests.

## Funding

This research was supported by the Extramural Research Programs of U.S. National Institutes of Health: NIMH (R01MH126518, RF1MH117053, R01MH111429, and S10MH124745), NINDS (R01NS091236 and RF1NS086085), NIAAA (P60AA011605), NICHD (P50HD103573), and OD (S10OD026796).

## Author contributions

V.M. and Y.-Y.I.S. conceived the project. T.-H.H.C. conducted the experiments. T.-A.E.N., B.W.L., N.K.B., and P.K.M. analyzed the data. T.-A.E.N. and V.M. wrote the manuscript with contributions from N.K.B. and input from all authors. V.M. and Y.-Y.I.S. supervised the study and provided funding support.

## Competing interests

The authors declare no competing interest.

## Data and materials availability

All original code used for the analysis and all data utilized in this study, will be deposited online and made available without restrictions.

## Supplementary Information

### I. Supplementary Methods

#### Animal preparation

Thy1-GCaMP6f transgenic male Long-Evans rats (RRID: RRRC_00830, Rat Resource & Research Center, Columbia, MO) weighing between 300 and 600 g were used in this study. These rats express the fluorescent calcium activity indicator, GCaMP, under Thy1 promoter, which allows for measurement of cortical output activity from pyramidal neurons. 8 rats in total were used for the free motion experiment. All procedures were conducted in accordance with the National Institutes of Health Guidelines for Animal Research (Guide for the Care and Use of Laboratory Animals) and approved by the University of North Carolina (UNC) Institutional Animal Care and Use Committee. The experimental designs followed the ARRIVE guidelines.

The rats were first anesthetized with 5% isoflurane and then maintained under anesthesia with a constant flow of 2-2.5% isoflurane mixed with medical air. Rectal temperature was continuously monitored and maintained within 37 ± 0.5 °C. The rats were placed on a stereotactic frame (Model 962, Kopf Instruments) with ear bars and a tooth bar. The scalp was removed to expose the skull (∼1×1 cm). Burr holes were prepared according to experimental coordinates. Specifically, the following coordinates were used: PrL (AP = 3.3 mm, ML = 0.8 mm, DV= 3.5 mm), Cg (AP = 1 mm, ML = 0.8 mm, DV = 2 mm), RSC (AP = -2.2 mm, ML = 0.7 mm, DV = 2 mm), and AI (AP = 3.2 mm, ML = 4.2 mm, DV = 3.5 mm).

Next, four miniature brass screws (Item #94070A031, McMaster Carr, Atlanta, GA) were anchored to the skull, and multimode optical fibers (200 μM core; NA: 0.37) were chronically implanted into the experimental coordinates. The surface of the skull was then covered with dental cement to seal the implanted components. Postoperative analgesics included Bupivacaine (2.5mg/ml subcutaneous), Lidocaine (2.5% topical) and meloxicam (1.5mg/kg), and Neomycin and polymyxin B sulfates and dexamethasone ophthalmic ointment. USP (Baushch & Lomb) was administered to prevent excessive dryness and infection of the eyes. All rats were allowed at least 1 week for recovery before any further experiments were conducted.

#### Multisite GCaMP recordings

We used a 4-channel fiber-photometry platform that was capable of spectrally resolved measurements (1-6). Excitation of the GCaMP signal was achieved through interleaved 488 nm and 405 nm diode lasers (OBIS Galaxy Laser 1236444 and 1236439, respectively) that provided calcium activity and motion correction reference signals, respectively. The lasers were launched into a dichroic mirror set (OBIS Galaxy Laser Beam Combiner), equally split into four outputs by a 1-to-4 fan-out fiber optic bundle (BF42LS01, Thorlabs), and delivered into a fluorescence cube (DFM1, Thorlabs). Additional neutral density filters (NEK01, Thorlabs) were placed to adjust the final laser power before entering the fluorescence cube. The fluorescence cube contained a dichroic mirror (ZT405/488/561/640rpcv2, Chroma Technology Corp) to reflect and launch the laser beam through an achromatic fiber port (PAFA-X-4-A, Thorlabs) into the core of a 105/125 mm core/cladding multimode optical fiber patch cable. The distal end of the patch cable was connected to an implantable optical fiber probe for both excitation laser delivery and emission fluorescence collection.

The fluorescence emitted from the fiber probe was collected and transmitted back along the patch cable, passed through the dichroic mirror and an emission filter (ZET405/488/561/640mv2, Chroma Technology Corp), then was launched through an aspheric fiber port (PAF-SMA-11-A, Thorlabs) into the core of an AR-coated 200/230 mm core/cladding multimode patch cable (M200L02S-A, Thorlabs). This cable was connected to a spectrometer (QE Pro-FL, Ocean Optics, Largo, FL) for spectral data acquisition, which was operated by a UI software OceanView (Ocean Optics, Largo, FL). We used a programmable Med-PC^®^ (Med Associates, Inc., Fairfax, VT) interface to synchronize the laser outputs, video recording, and spectrometer recordings with TTL pulses. The interleaved laser outputs were applied at a 50% duty cycle with 50 ms pulse duration. Emission spectra by the interleaved 488 and 405 nm lasers were collected at 1 ms after each laser switching with a 25 ms sampling window, resulting in a 10 Hz effective sampling rate for both GCaMP signal time-course derived from 488 and 405 nm excitation. The video was collected at a frame rate of 10 Hz and synchronized with 488 laser triggers.

Prior to attaching the fiber-photometry patch cables to the implanted fiber ferrules on each rat, background spectra were collected for the 405 and 408 nm lasers. This was achieved by directing the patch cable fiber tips towards a nonreflective surface in the fiber-photometry recording room. The resulting background spectra were subsequently subtracted during the data analysis process. To avoid animal stress during attaching the four patch cables to the four implanted fiber ferrules on each rat, the rats were initially anesthetized by 5% isoflurane, then maintained under anesthesia by a constant flow of 2-2.5% isoflurane mixed with medical air during attaching the patch cables. After the patch cables were attached, the rat was returned to its home cage (which measured 37 cm x 33.5 cm in floor size and had a height of 18 cm), and allowed to recover from anesthesia for 30 min before fiber-photometry recording. We placed the rodents in their familiar home cage environment during the experiment to capture neural dynamics with minimal influence from novel environmental stimuli, and to reduce potential influences of stress and anxiety levels.

#### GCaMP data preprocessing

GCaMP data preprocessing steps were the same as described in a previous study (1). To quantify GCaMP neural activity, the recorded signal *Y*(*t*)^(*s*)^ in each rodent *s* at each acquisition time point *t* was decomposed across light wavelengths ranging from and 500 nm to 600 nm and fitted with the model:

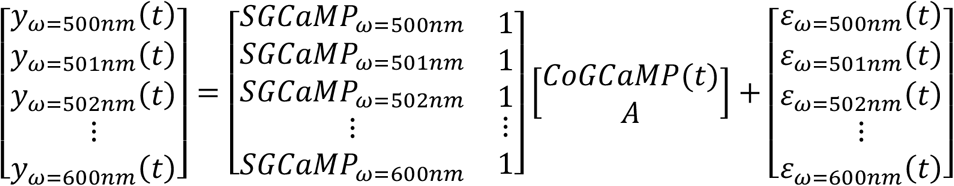

where *S*_*GCaMP*_ is the normalized reference emission spectrum of GCaMP across light wavelengths, *Co*_*GCaMP*_(*t*)^(*s*)^ is the estimated coefficient of interest in animal *s* at time *t, A*^(*s*)^ is the mean or DC offset of signal *Y*(*t*)^(*s*)^over time in *s*, and *ε*(*t*)^(*s*)^ is the residual in animal *s* at time *t*. The derived *Co*_*GCaMP*_(*t*)^(*s*)^ was then detrended and mean-centered. To measure the time-frequency energy distributions of GCaMP activity, *Co*_*GCaMP*_(*t*)^(*s*)^ was decomposed into a time-frequency function using the continuous wavelet transformation with complex Morlet wavelets as the mother wavelet. On the basis of the energy distributions of GCaMP activity, the GCaMP signals were band-pass–filtered at 0.1 to 1.5 Hz for further analyses.

#### Functional connectivity analysis and hierarchical clustering of RSC, Cg, PrL, and AI GCaMP signals during spatial exploration

Functional connectivity between regions was computed using Pearson correlation between the GCaMP signals (over time) in each pair of regions for each animal. For each region pair, we determined whether the distribution of the correlations across animals was zero. Significance testing was performed using a two-sided Wilcoxon signed-rank test (*n* = 8 rats), with FDR correction for across six comparisons. Then, significance testing was performed over comparisons between all region pairs with a paired Wilcoxon signed-rank test (*n* = 8 rats), with FDR correction for across all 15 possible combinations of region pairs. We required *P* < 0.05 for significance. Hierarchical clustering with complete linkage (7) was then performed, using one minus the mean r across animals as a metric of distance between nodes.

#### Phase lag index (PLI) analysis of RSC, Cg, PrL, and AI GCaMP signals during spatial exploration

We examined phase relations between GCaMP signals using the phase lag index (PLI). The PLI is the mean sign of the phase difference in a pair of signals. For a time-series of length T from regions i and j in animal s,

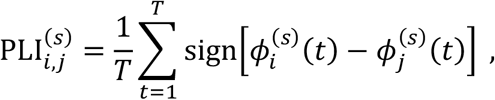

where 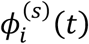 and 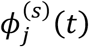 are the phases of the Hilbert-transformed GCaMP signals from animal *s*, at time *t*, and in regions *i* and *j* respectively. If the PLI is positive, region *i* precedes region *j* in phase, and if the PLI is negative, *i* lags *j*. If the PLI is zero, the regions are either perfectly synchronized or not presenting any systematic phase relation. For each pair of regions, we determined whether their PLI is significantly different from zero, using a two-sided Wilcoxon signed-rank test (*n* = 8 rats). We FDR corrected the *P* values of these tests for 6 comparisons and required *P* < 0.05 for significance.

#### MVAR modeling of directed regional interactions during spatial exploration

To investigate how GCaMP signals from different regions mutually influence each another, we fit a multivariate autoregression model (MVAR) model. This model allowed us to infer linear directional weights between our neural recordings:

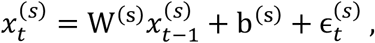

where, for animal *s*, 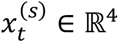 consists of the GCaMP signals in RSC, Cg, PrL, and AI at time *t, W*^(*s*)^ ∈ ℝ^4×4^ is a directional weight matrix, *b*^(*s*)^ ∈ ℝ^4^ is a bias, and 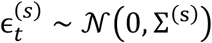 is a Gaussian random variable with Σ^(s)^ ∈ ℝ^4×4^ a covariance matrix. Furthermore, letting *T* be the length of the time series, 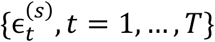 is a white-noise process (*T* = 11800). For region *i*, the outflow of weights is 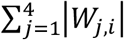. For each region pair, we tested whether the difference between their outflow of weights is zero with a two-sided Wilcoxon signed-rank test (*n* = 8 rats). We FDR corrected the *P* values of these tests for 6 comparisons and required *P* < 0.05 for significance.

#### Estimation of the kinematics of spatial exploration from video recordings

To estimate kinematic measures from video recordings, we extracted the trajectory through time of the brightly lit cap on the rodent’s head to which the optical fiber was connected. To detect the position of the cap in each video frame, we computed a histogram of the luminosity across pixels in that frame and identified the local maximum in that histogram that was largest in luminosity. Luminosities above the local minimum immediately preceding that local maximum were inferred to be from pixels of the cap, while luminosities below it were inferred to be from other parts of the image. Among the pixels of higher luminosity than the local minimum, the center of mass was estimated and taken to be the position of the animal’s head. The center of mass was defined as the mean of the coordinates of the cap pixels weighted by their luminosities. If no local maximum in the luminosity histogram was observed in a frame, such as when the cap was not present in the frame (in case the rat’s head wandered out of frame or was hidden under its body), the corresponding time point was excluded from further analysis. Once the 2-dimensional coordinates of the cap position in each frame were identified, the linear velocity, angular velocity, and linear acceleration across three consecutive time points were computed. For rat *s* at time *t*, linear velocity is given by:

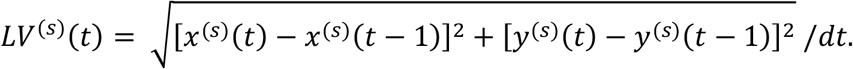

Angular velocity is given by:

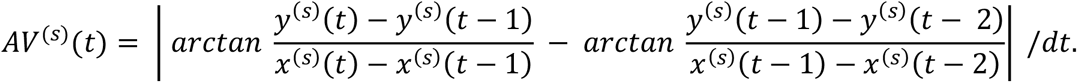

And linear acceleration is given by:

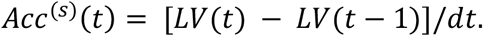

#### Relation between GCaMP neuronal activity and kinematics during spatial exploration

For each animal, we computed the Pearson correlation between each kinematic measure (linear velocity, angular velocity, or acceleration), and the GCaMP signal in each of the four brain regions. For each kinematic measure, we then determined whether the correlation was different from zero, using a two-tailed Wilcoxon signed-rank test. We FDR corrected the P values of these tests for each kinematic measure across 4 comparisons and required P<0.05 for significance.

#### State-space modeling of neural dynamics during spatial exploration

We applied a Bayesian switching linear dynamical system (BSDS) model (8), as in our previous studies (1). BSDS describes time-series data using continuous latent variables endowed with switching, autoregressive linear dynamics. The switching of dynamics is governed by discrete latent variables, which evolve as a Markov chain. We learned this model’s parameters using exact coordinate ascent variational inference (35).

We now describe BSDS’s probabilistic model. For subject *s* and a time-series of length *T*, the observations are 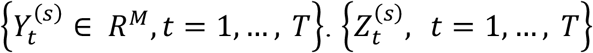 is a discrete, latent, homogenous Markov chain with state space {1, …, *K*}. 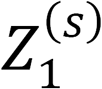 ∼ Categorical(*π*), 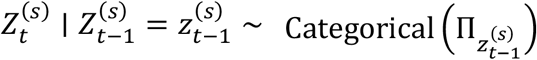, where *π* is a *K* – 1 probability simplex and ∏ is a row stochastic matrix.

Then, for 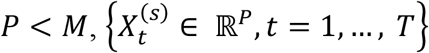 is a latent vector autoregressive first-order Gaussian process, such that

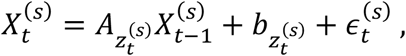

where 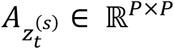 is a dynamics matrix, 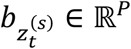 is a bias, and 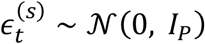 is a Gaussian random variable with *I*_*P*_ ∈ ℝ^*P*×*P*^ an identity matrix. Finally, the observations

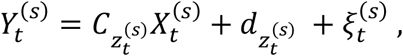

where 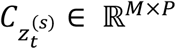 is an emissions matrix, 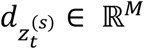 is an emissions bias, and 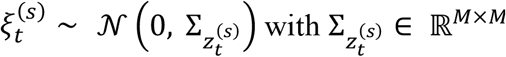 a covariance matrix. Priors for the *A*’s and *b*’s implement automatic relevance determination, and, in general, priors are chosen to enable inference using exact coordinate-ascent mean-field variational inference. More detail is provided in (35).

We applied BSDS to the GCaMP recordings from RSC, Cg, PrL, and AI (*M* = 4, *T* = 11800). We set the maximum dimensionality of the discrete latent process *K* = 10 and the dimensionality of the continuous latent process to *P* = 3, and performed inference jointly on the 8 animals. We emphasize that BSDS was learned exclusively from our neural data (i.e., without any behavioral data). We then ran the Viterbi algorithm to identify the most likely sequence of the latent discrete states, the Viterbi path, for each animal. 7 out of the 10 possible states appeared in these most likely sequences of discrete states. We were interested here in the discrete states: they identify spatiotemporal dynamics common through each neural recording session and over all rodents, and they provide for the segmentation of our complex time series into syllables of neural activity (i.e., brain states).

#### Occupancy rate and mean lifetime of latent brain states

Occupancy rate and mean lifetime of latent brain states We defined the occupancy rate for each animal and each brain state as the number of occurrences of the state in the Viterbi path divided by the length of the time series. We defined the mean lifetime as the mean duration in time of consecutive appearances of the state in the animal’s Viterbi path.

#### Distribution of kinematic measures over latent brain states

To determine whether linear velocity, angular velocity, and acceleration varied across the latent brain states uncovered by BSDS, we computed, for each animal and latent state, the mean of each kinematic metric in that state and animal. We first tested whether all the latent states have the same distribution of linear velocity using the Friedman test (k=7 states, n=8 rats). P values from the Friedman tests were computed using exact null distributions (8). We required P<0.05 for significance. Similar analyses were conducted using angular velocity and acceleration.

#### Relation between brain state-specific neural activity and time-varying kinematics during spatial exploration

To evaluate whether GCaMP activity in the brain regions covaried with kinematic measures across latent brain states, we first extracted the time points of the GCaMP signals associated with each brain states identified by BSDS. To do so, we used the following procedures. For array *x* and Boolean array *y* of the same length as *x*, let [*x* | *y*] denote an array formed from indexing *x* by exactly the indices where *y* is true, and, for *a, b* ∈ ℝ, let *I*_*a*_(*x*) be the array obtained by applying

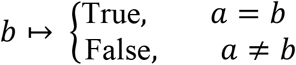

elementwise over *x*. For brain region *i*, locomotion measure *j*, and animal *s*, let

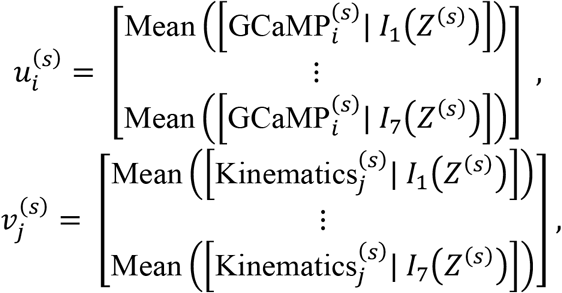

where, Mean(*u*) denotes the mean of *u*, 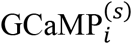 is the time-series array of GCaMP signal in brain region *i* in animal *s*, 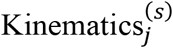 is the time-series array of locomotion measure *j* in animal *s*, and *Z*^(*s*)^ is the Viterbi path in animal *s*. We then computed 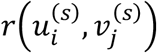 for each *i, j*, and *s*, where *r* (*u, v*) is the Pearson correlation coefficient between *u* and *v*.

For each *i* and *j*, we determined whether the correlation between mean brain state-specific GCaMP activity in region *i* and kinematic measure *j* was different from zero correlation across the states, using a two-sided Wilcoxon signed-rank test (*n* = 8 rats). Then, for each DMN node, we determined whether the correlation between mean GCaMP activity in the DMN node and linear velocity was higher than the correlation between mean GCaMP activity in the AI and linear velocity. We used a two-sided Wilcoxon signed-rank test (*n* = 8 rats) and FDR corrected the *P* values across 3 comparisons. For both the tests of the correlations and their differences, we required *P* < 0.05 for significance.

#### Relation between brain state-specific spectral entropy of neural activity and time-varying kinematic measures during spatial exploration

To investigate how spectral entropy covaried with kinematic metrics across latent brain states, we extracted time points belonging to each latent state using the following procedure. Continuing with the notation from the previous section, for brain region *i* and animal *s*, let

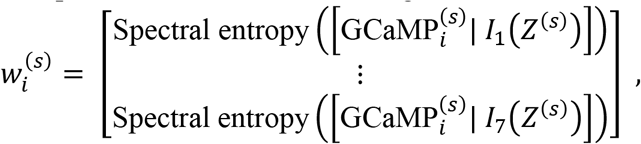

where Spectral entropy(*u*) was calculated in the following way.

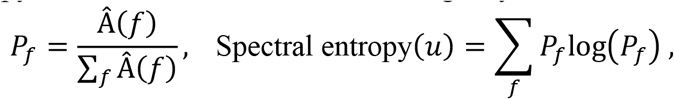

where *f* is a frequency bin and Â(*f*) is the amplitude of the power spectral density of *u*, computed with a Welch transform, at *f*. Spectral entropy was computed the arrays 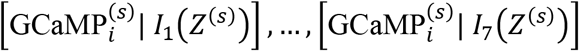, where power spectra were computed with a frequency binning of 0.04 Hz. We then computed 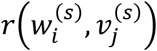 for each *i, j*, and *s*. For each *i* and *j*, we tested whether the spectral entropy of GCaMP activity in region *i* and locomotion measure *j* have zero correlation over the states with a two-sided Wilcoxon signed-rank test (*n* = 8 rats). We required *P* < 0.05 for significance.

#### Relation between brain state-specific functional connectivity and time-varying kinematics during spatial exploration

To investigate whether brain state-specific functional connectivity covaried with kinematic metrics across latent brain states, we used the following procedure. Again, continuing with the notation from the previous section, for brain region pair (*k, l*) and animal *s*, let

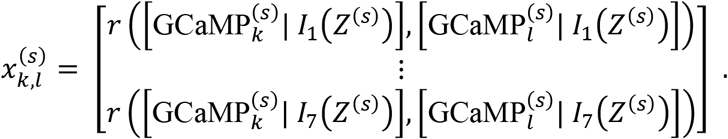

We then computed 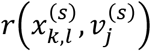 for each *i*, (*k, l*), and *s*. For each *i* and (*k, l*), we determined whether the correlation between functional connectivity of region pair (*k, l*) and kinematic measure *j* was significantly different from zero across the brain states, using a two-sided Wilcoxon signed-rank test (*n* = 8 rats). We required *P* < 0.05 for significance.

### II. Supplementary Results

#### Neural activity in the DMN encodes time-varying angular velocity and acceleration during spatial exploration

We investigated how kinematic measures covaried with GCaMP neural activity across brain states (**Figure S1**). Angular velocity was negatively correlated with mean GCaMP activity in the 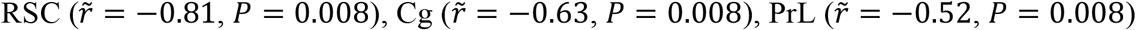, and 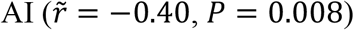 (**Figure S1A**). The strength of the negative correlation between angular velocity and mean GCaMP signal was higher in the RSC than in the AI (FDR corrected *P* = 0.023) (**Figure S1B**). Similarly, acceleration was negatively correlated with mean activity in the 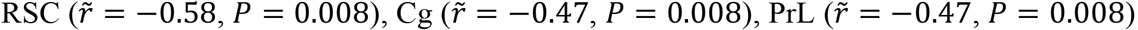, and 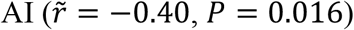, (**Figure S1C**). No significant differences were found between the regions in the strength of the correlations between acceleration and mean GCaMP signal (**Figure S1D**). These results demonstrate a pattern convergent with those observed with linear velocity.

#### Spectral entropy of neural activity in the DMN changes with angular velocity and acceleration during spatial exploration

We investigated how kinematic measures covaried with spectral entropy of GCaMP neural activity across brain states (**Figure S2**). Angular velocity was positively correlated with spectral entropy in the 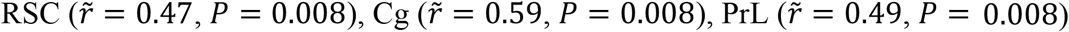, and 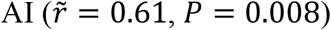 (**Figure S2B**). Similarly, acceleration was positively correlated with spectral entropy in the 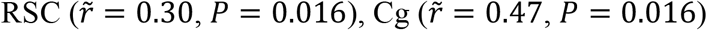, and 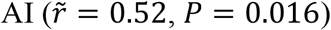, but not significantly in the 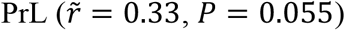 (**Figure S2C**). These results demonstrate a pattern convergent with those observed with linear velocity.

#### Functional connectivity of DMN nodes encodes time-varying angular velocity and acceleration during spatial exploration

We investigated how kinematic measures covary with functional connectivity across brain states (**Figure S3 and S4**). Angular velocity was negatively correlated with functional connectivity between the RSC and 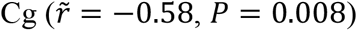 (**Figure S3A**) and between the Cg and 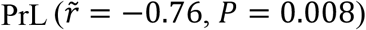 (**Figure S3A**). Angular velocity was positively correlated with functional connectivity between the RSC and 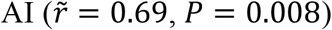 (**Figure S3B**). Acceleration was negatively correlated with functional connectivity between the RSC and 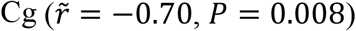 (**Figure S4A**) and positively correlated with functional connectivity between the RSC and 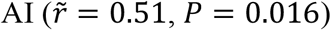 (**Figure S4B**). These results demonstrate a pattern convergent with those observed with linear velocity.

### III. Supplementary Figures

**Figure S1.**
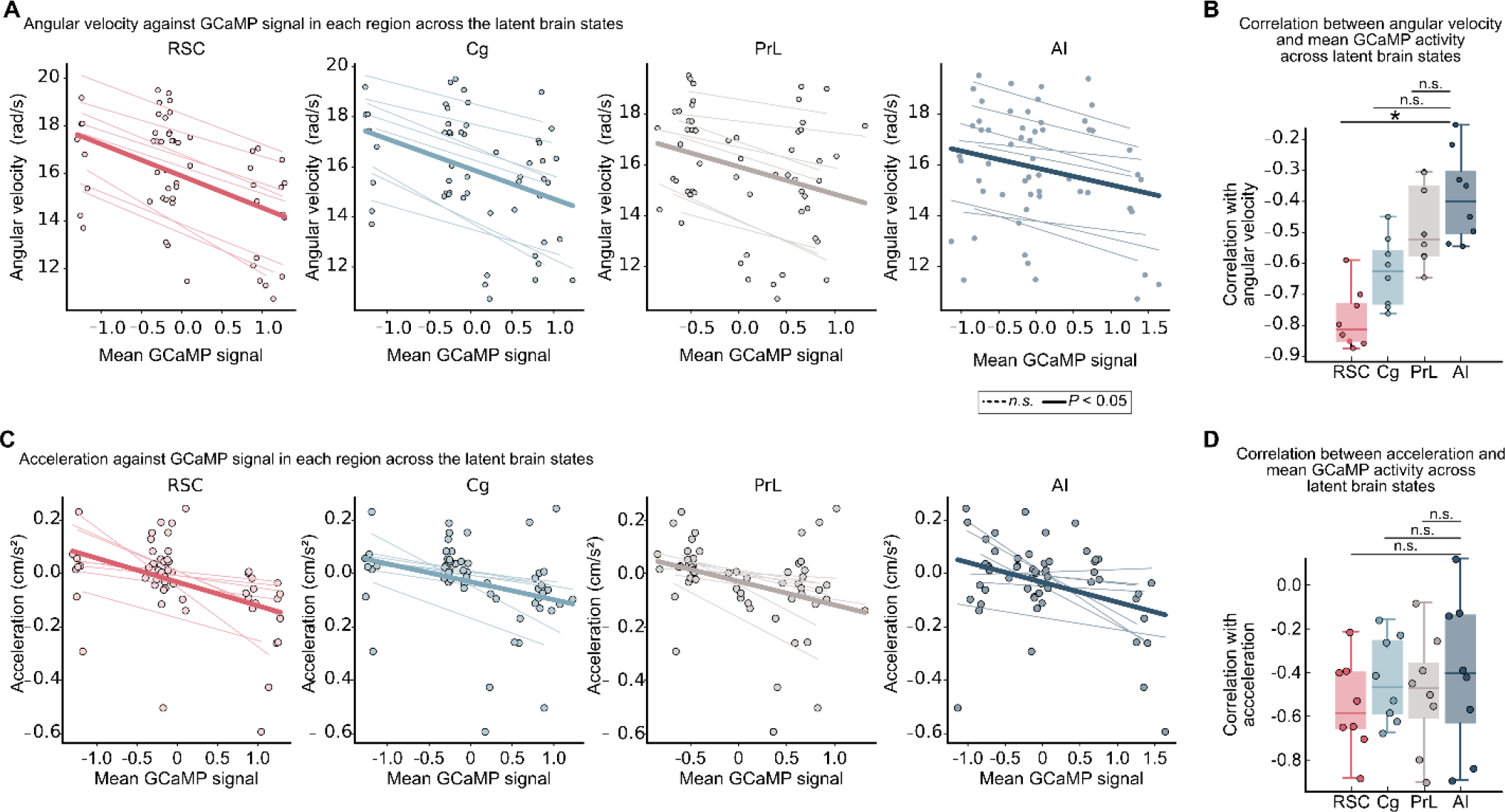
DMN activity encodes time-varying kinematics during spatial exploration. **(A)** Angular velocity as a function of mean GCaMP signal amplitude in the RSC, Cg, PrL, and AI across latent brain states. Angular velocity was negatively correlated with mean GCaMP activity in the RSC, Cg, PrL, and AI. Lighter lines represent linear fits across latent brain states in each rat, and each circle represents data from a specific rat and latent brain state. Dark lines represent linear models across all rodents (n.s. *P* ≥ 0.05). **(B)** Strength of correlation between angular velocity and mean GCaMP signal. The magnitude of the negative correlation was greater in the RSC than in the AI (* FDR corrected *P* < 0.05, n.s. FDR corrected *P* ≥ 0.05). **(C)** Acceleration as a function of mean GCaMP signal amplitude in the RSC, Cg, PrL, and AI across latent brain states. Acceleration was negatively correlated with mean GCaMP activity in the RSC, Cg, PrL, and AI. Lighter lines represent linear fits across latent brain states in each rat, and each circle represents data from a specific rat and latent brain state. Dark lines represent linear models across all rodents (n.s. *P* ≥ 0.05). **(D)** Strength of correlations between acceleration and mean GCaMP signal. No significant differences were found between the regions in the strength of the correlations between acceleration and mean GCaMP signal (n.s. FDR corrected *P* ≥ 0.05). Box and whisker plots show 25^th^ and 75^th^ percentiles, and horizontal lines represent median values. Dots in each figure represent data points from individual rats.

**Figure S2.**
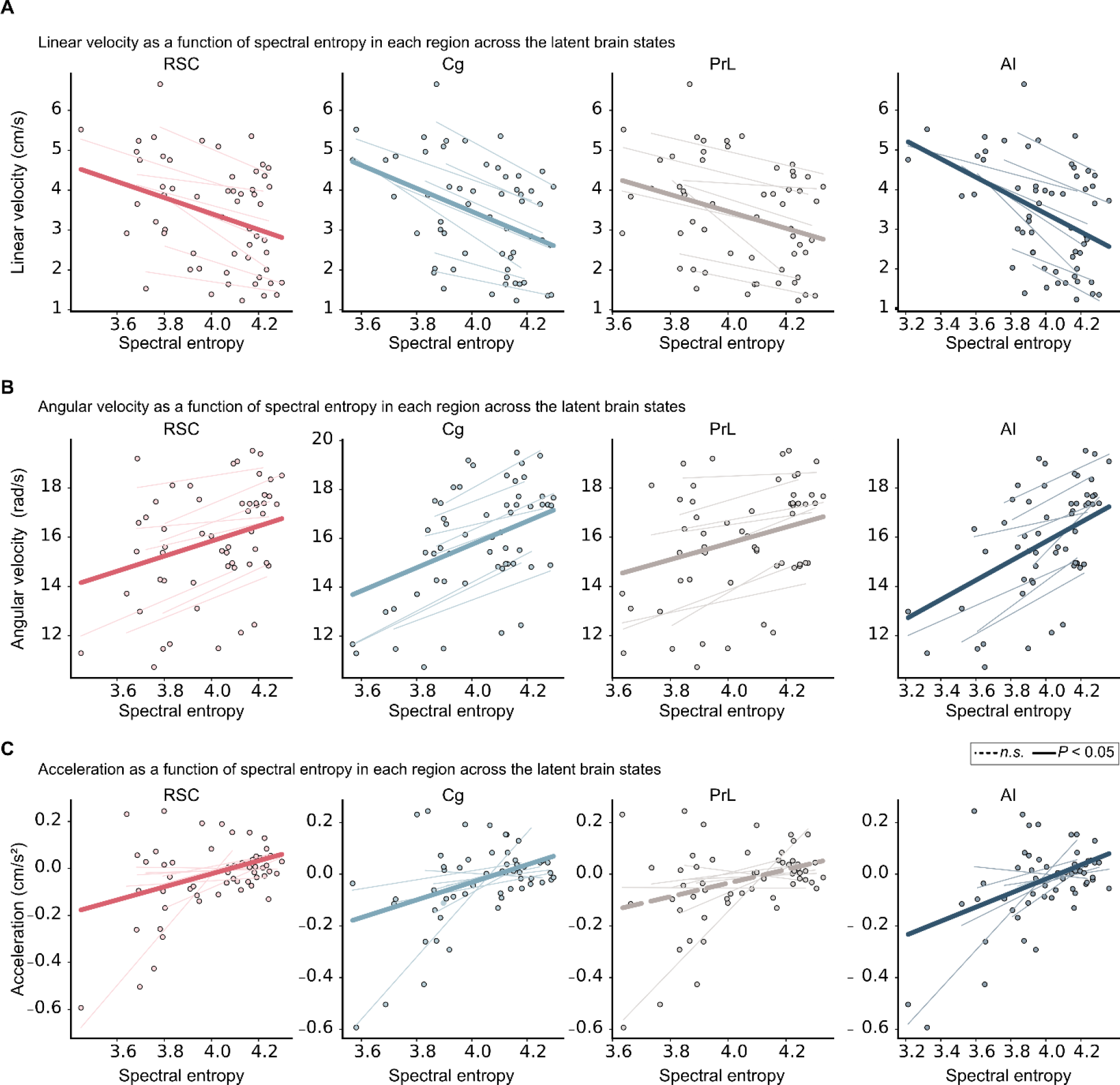
Spectral entropy changes with time-varying kinematics during spatial exploration. **(A)** Linear velocity, **(B)** Angular velocity, and **(C)** Acceleration as a function of spectral entropy in the RSC, Cg, PrL, and AI, across latent brain states. Linear velocity was negatively correlated with spectral entropy in each region. Angular velocity was positively correlated with spectral entropy in each region, and acceleration was also positively correlated with spectral entropy in the RSC, Cg, and AI, though not significantly in the PrL. Lighter lines represent linear fits across latent brain states in each rat, and each circle represents data from a specific rat and brain state. Dark lines represent linear models across all rodents (n.s. *P* ≥ 0.05).

**Figure S3.**
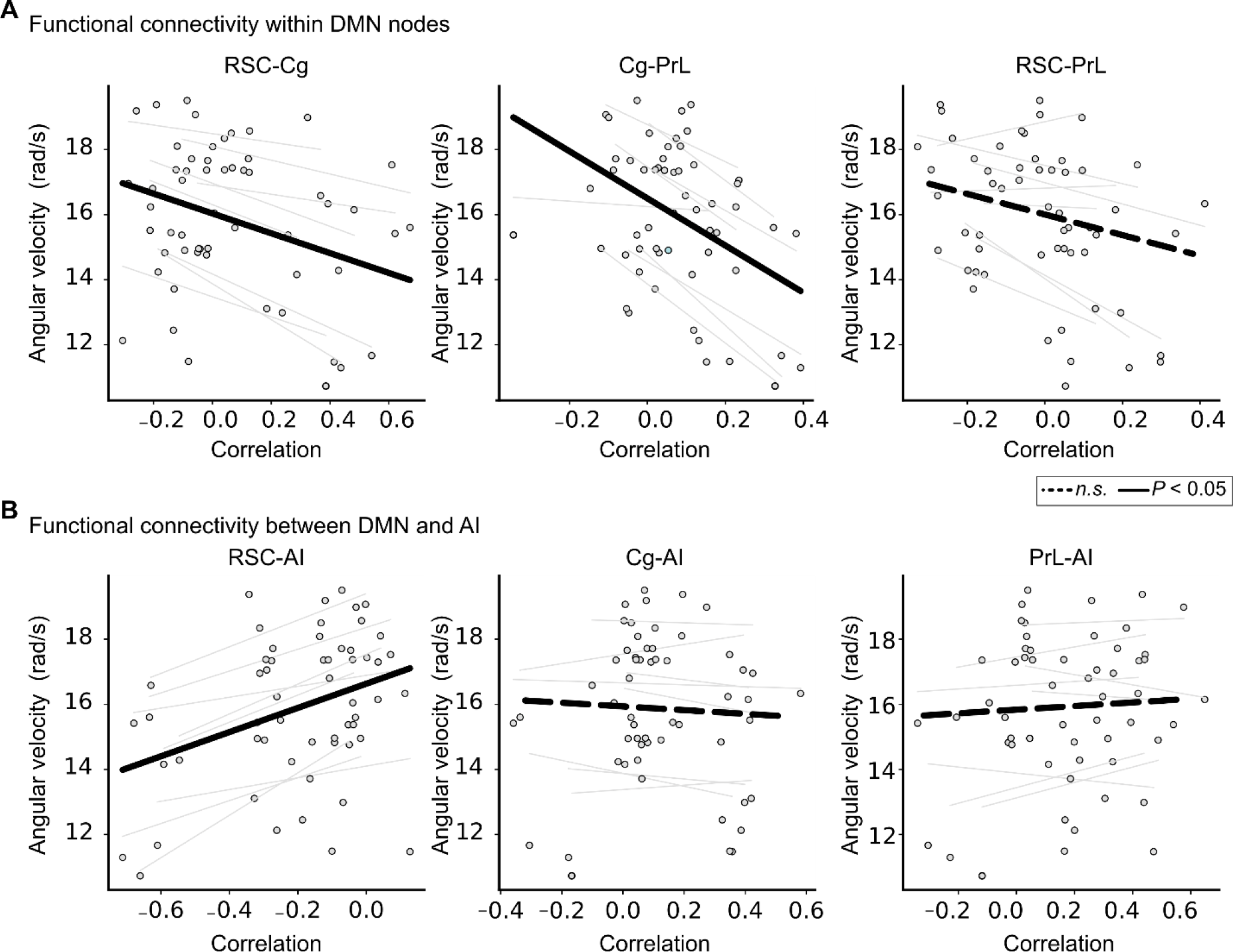
DMN connectivity encodes time-varying angular velocity during spatial exploration. **(A)** Angular velocity as a function of intra-DMN functional connectivity between the RSC, Cg, and PrL, across latent brain states. Angular velocity was negatively correlated with functional connectivity between the RSC and Cg and between the Cg and PrL. **(B)** Angular velocity as a function of connectivity between the AI node of the salience network and the RSC, Cg, and PrL nodes of the DMN. Angular velocity was positively correlated with functional connectivity between the RSC and AI. Lighter lines represent linear fits across brain states in each rat, and each circle represents data from a specific rat and latent brain state. Dark lines represent linear models across all rodents (n.s. *P* ≥ 0.05).

**Figure S4.**
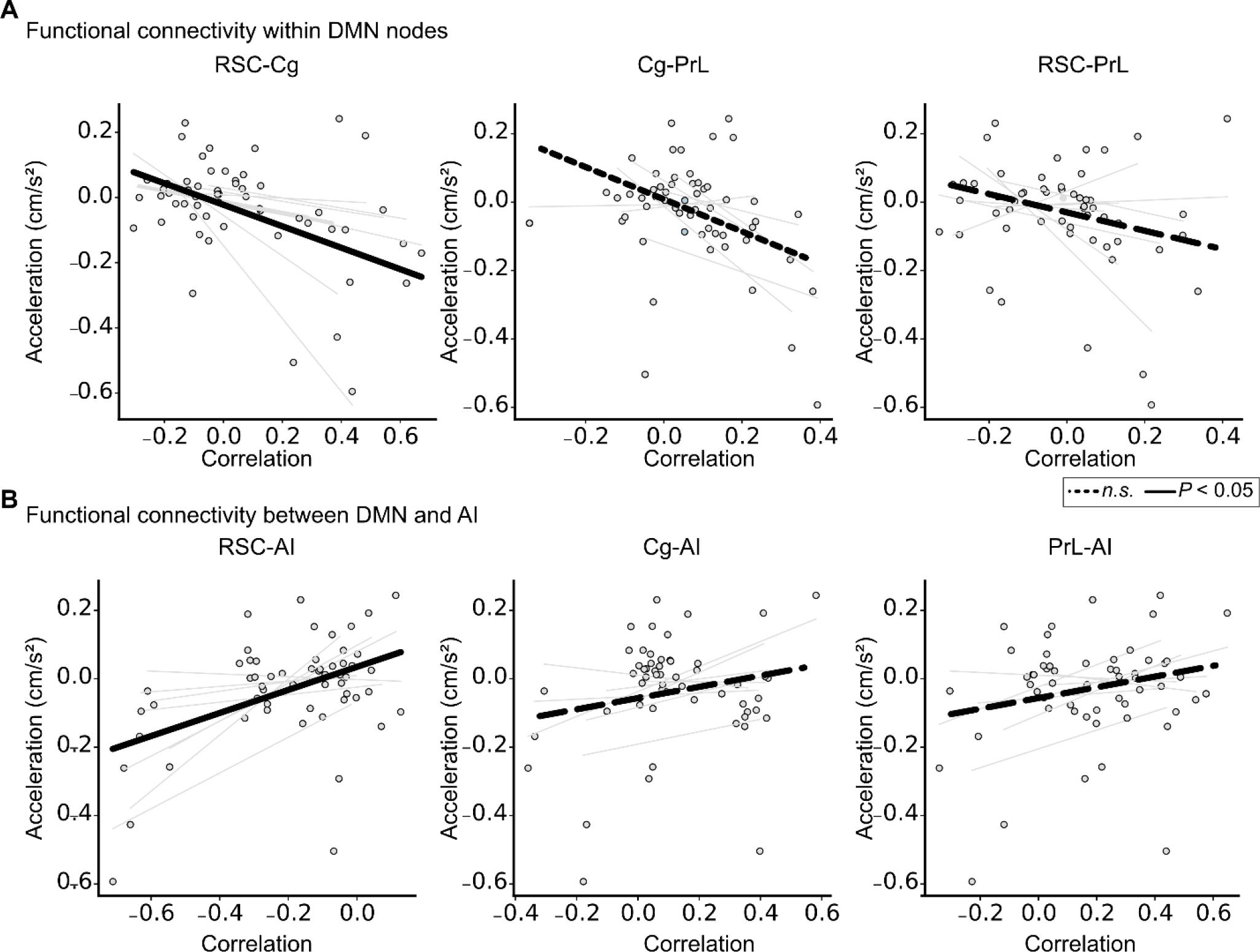
DMN connectivity encodes time-varying acceleration during spatial exploration. **(A)** Acceleration as a function of intra-DMN functional connectivity between the RSC, Cg, and PrL, across latent brain states. Acceleration was negatively correlated with functional connectivity between the RSC and Cg. **(B)** Acceleration as a function of connectivity between the AI node of the salience network and the RSC, Cg and PrL nodes of the DMN. Acceleration was positively correlated with functional connectivity between the RSC and AI. Lighter lines represent linear fits across brain states in each rat, and each circle represents data from a specific rat and latent brain state. Dark lines represent linear models across all rodents (n.s. *P* ≥ 0.05).

## Notes

### Competing Interest Statement

The authors have declared no competing interest.

